# The atmospheric pressure capillary plasma jet is well-suited to supply H_2_O_2_ for plasma-driven biocatalysis

**DOI:** 10.1101/2025.01.07.631711

**Authors:** Tim Dirks, Davina Stoesser, Steffen Schüttler, Frank Hollmann, Judith Golda, Julia E. Bandow

## Abstract

Plasma-generated H_2_O_2_ can be used to fuel biocatalytic reactions that require H_2_O_2_ as co-substrate such as the conversion of ethylbenzene to (*R*)-1-phenylethanol ((*R*)-1-PhOl) catalyzed by unspecific peroxygenase from *Agrocybe aegerita* (r*Aae*UPO). Immobilization was recently shown to protect biocatalysts from inactivation by highly reactive plasma-produced species, however, H_2_O_2_ supply by the employed plasma sources (µAPPJ and DBD) was limiting for r*Aae*UPO performance. In this study we evaluated a recently introduced capillary plasma jet for suitability to supply H_2_O_2_ *in situ*. H_2_O_2_ production was modulated by varying the water concentration in the feed gas, providing a greater operating window for applications in plasma-driven biocatalysis. In a static system after 80 min of biocatalysis, a turnover number of 44,199 mol_(*R*)-1-PhOl_ mol^-1^_r*Aae*UPO_ was achieved without significant enzyme inactivation. By exchanging the reaction solution every 5 min, a total product yield of 122 µmol (*R*)-1-PhOl was achieved in 700 min run time, resulting in a total turnover number of 174,209 mol_(*R*)-1-PhOl_ mol^-1^_r*Aae*UPO_. We conclude that the capillary plasma jet, due to its flexibility regarding feed gas, admixtures, and power input, is well-suited for *in situ* H_2_O_2_ generation for plasma-driven biocatalysis tailoring to enzymes with high H_2_O_2_ turnover.

## Introduction

Peroxygenases use hydrogen peroxide to carry out oxyfunctionalization reactions such as stereoselective hydroxylations [1] and epoxidations [2,3], which are chemically difficult (or impossible) to achieve, making them appealing enzymes for biocatalysis [4]. However, their industrial application is limited since they suffer from hydrogen peroxide-induced inactivation upon disruption of the heme cofactor [5,6]. H_2_O_2_ levels need to be tightly controlled to not inactivate the biocatalyst. To this end, recently, numerous approaches for *in situ* H_2_O_2_ generation have been tested, such as electrochemical [7,8], photocatalytic [9,10], and thermoelectric methods [11] or the use of enzyme cascades [1]. Another approach for *in situ* H_2_O_2_ generation is the utilization of atmospheric pressure plasmas. These plasmas generate, among many other species, H_2_O_2_ as a long-living species [12]. A first proof-of-principle study using a dielectric barrier discharge (PlasmaDerm, Cinogy, Germany) as plasma source revealed that plasma-generated H_2_O_2_ can be used to drive the reaction of H_2_O_2_-utilizing enzymes [13]. This initial study revealed two major limitations, namely enzyme stability under plasma conditions and low H_2_O_2_ supply by that plasma source. Enzyme inactivation is a well-known effect of plasma treatments [14–16]. We would like to highlight three mechanisms: (i) inactivation of cofactors such as heme [5,6], (ii) the modification of amino acid side chains that are involved in catalysis or are crucial for the enzyme structure (especially histidines or cysteines) [15,17], and (iii) the degradation of enzymes by cleavage of peptide bonds, which is presumably induced especially by short-living species [14]. To protect enzymes under plasma treatment conditions, immobilization was identified as a promising tool [18]. By immobilizing enzymes on inert carrier materials, an enzyme-free buffer zone is created between the enzyme-loaded beads and the plasma-exposed liquid surface, allowing the highly reactive short-living species to recombine before they could interact with the enzymes. However, the functional groups used to immobilize enzymes on the carrier material have a profound impact on the plasma stability of the biocatalyst [18, 19]. In addition to protecting against plasma-induced inactivation, immobilization allows to reuse the enzymes, increasing the overall cost efficiency of the process. To increase H_2_O_2_ production the atmospheric pressure plasma jet (µAPPJ) was evaluated as an alternative plasma source [20]. Higher H_2_O_2_ production was achieved, the treatable volume was increased, and coupling to online biocatalysis was possible. However, 25% of the feed gas could be saturated with water since higher admixtures led to the extinction of the plasma. The input power of the system could not be increased beyond 1.5 - 2 W, since increasing arc discharges could occur that damage the electrodes. The H_2_O concentration in the feed gas is an important parameter for plasma-induced H_2_O_2_ generation [21-25], and H_2_O_2_ production with the µAPPJ was limited. Thus, In the present study, the capillary plasma jet (CPJ) was used, which allows higher input power and thus higher H_2_O concentrations in the feed gas [26] (Figure 1). We investigated H_2_O_2_ formation and biocatalysis performance with the recombinantly produced unspecific peroxygenase from *Agrocybe aegerita* (r*Aae*UPO) [27]. Furthermore, we tested alternatives for the amino-functionalized ReliZyme beads (HA403 M) with regard to their utility for plasma-driven biocatalysis.

**Figure 1:**
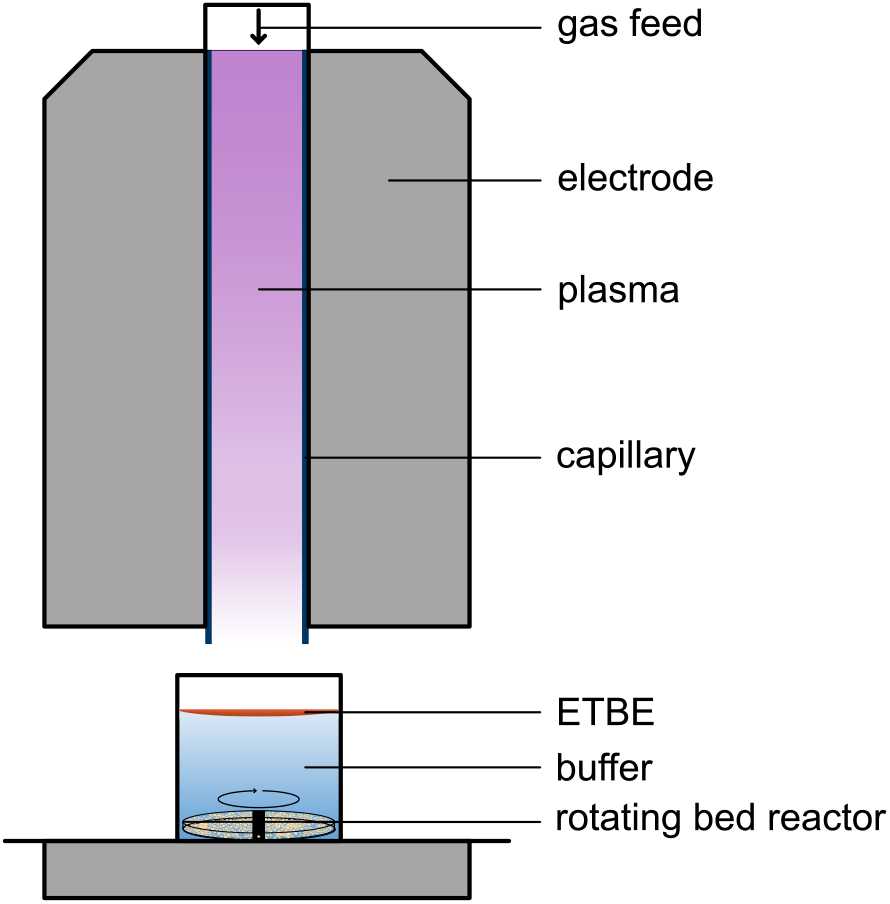
Plasma-driven biocatalysis setup used in this study. The feed gas of the capillary jet was routed through a water-containing bubbler and subsequently guided through a capillary surrounded by two plane-parallel electrodes. The sample, potassium phosphate buffer (5 ml, 100 mM, pH 7) containing 50 mM of the substrate ETBE, is exposed to the plasma effluent exiting the capillary. The rotating bed reactor containing the protein-loaded beads is located at the bottom of the vessel. Image was modified from [13,20,26].

## Experimental procedures

### Plasma source

The capillary plasma jet used in this study was identified as promising source for H_2_O_2_ generation and H_2_O_2_ transfer into liquid samples [28]. The radio frequency (RF)-driven (13.56 MHz) atmospheric pressure capillary plasma jet was used as described before [23] but with inner capillary (Hilgenberg, Germany) dimensions of 4.56 mm × 0.88 mm × 100 mm. The electrodes (itec Automation & Laser AG, Germany) had a width and length of 4 mm and 40 mm, respectively, resulting in a plasma volume of 4 mm × 0.88 mm × 40 mm. The wall thickness of the capillary was 0.22 mm, leading to a distance of 1.32 mm between the electrodes. Plasma was operated at a plasma power of (6.0 ± 0.6) W. The gas flow (2 slm He) was split and partially routed through a bubbler with cooled deionized water to enrich the feed gas with water molecules. The distance between the nozzle of the capillary jet and the sample was approx. 16 mm and the plasma zone ended 10 mm in front of the nozzle exit of the capillary.

### H_2_O_2_ measurement

H_2_O_2_ produced in the plasma-treated liquid was measured as described elsewhere [20]. In short, 5 ml potassium phosphate buffer (100 mM, pH 7) were exposed to plasma for different treatment times. Samples were diluted to appropriate concentrations using deionized water and mixed with reagents 1 and 2 of the commercially available Spectroquant H_2_O_2_ kit (Merck, Germany). After 5 min incubation, absorption was measured at 455 nm. H_2_O_2_ concentrations were calculated based on a calibration curve.

### Enzyme preparation and immobilization

r*Aae*UPO was purified as described previously [29]. For immobilization, two different types of non-directional carriers were used, namely amino (ECR8309F) and epoxy-butyl (ECR8285) functionalized Lifetech ECR resins (Purolite, Llantrisant, Wales) or amino (HA403 M) and ethyl-amino (EA403 M) functionalized beads (Resindion, Binasco, Italy). Beads (500 mg) were weight in a suitable vessel and washed thrice with 5 ml potassium phosphate buffer (100 mM, pH 7). The amino functionalized carrier materials were incubated in phosphate buffer containing 0.5 % (w/v) glutaraldehyde. After 3 h of incubation, beads were washed thrice with potassium phosphate buffer. Enzyme (2 nmol) was added, and immobilization was allowed to proceed overnight at 8°C with tubes shaking upside down.

### Binding efficiency

After immobilization, binding efficiency was determined measuring the remaining enzyme activity in the supernatant using 2,2’-azino-bis(3-ethylbenzothiazoline-6-sulfonic acid) (ABTS) as substrate. r*Aae*UPO (2 nmol) or supernatant was added to 100 mM sodium citrate buffer (pH 5) containing ABTS (5 mM). To start the reaction, an equivalent volume of H_2_O_2_ (2 mM) was added and product formation was monitored at 405 nm using a microplate reader (Biotek Epoch, Germany). Final concentrations were 2.5 mM ABTS, 1 mM H_2_O_2_, and 50 mM citrate. Unimmobilized r*Aae*UPO served as control.

### Biocatalysis setup

Plasma-driven biocatalysis was performed as described previously [20]. In short, 150 mg protein-loaded beads were transferred to a rotating bed reactor (build in-house, dimensions: ∅ 2 cm × 0.7 cm). The reactor was placed in a vessel filled with 5 ml potassium phosphate buffer (100 mM, pH 7) containing 50 mM ethylbenzene (ETBE). Plasma treatment was performed for up to 40 min, while every 5 min 150 µl sample were withdrawn for gas chromatography-based quantification of product as described elsewhere [13,20]. After biocatalysis, residual activities of protein-loaded beads were determined. To this end, beads were recovered from the rotating bed reactor and washed thrice with potassium phosphate buffer. Enzyme activity was determined as described for the unimmobilized enzyme but in a total volume of 1 ml. To ensure sufficient substrate supply, samples were incubated shaking during turnover. For a total of ten minutes reaction time, every two minutes aliquots of 100 µl were removed and measured at a wavelength of 405 nm using a microplate reader (Biotek Epoch, Germany). Enzyme activity was calculated based on the linear slope of the kinetic.

### Determination of the total turnover number (TTN)

For the long-term biocatalysis, 400 mg protein-loaded beads were transferred to a rotating bed reactor. To avoid H_2_O_2_ accumulation and product inhibition, the entire reaction solution was withdrawn and replaced with fresh solution every 5 or 10 min as indicated. Product formation was analyzed by GC. After addition of the fresh potassium phosphate ETBE mixture, plasma treatment was continued. Residual enzyme activity was measured at various time points and plasma treatment was applied until product formation per minute was below 0.05 µmol min^-1^.

## Results and Discussion

Technical, non-thermal atmospheric pressure plasmas can be used as the H_2_O_2_ producing source to drive the enzymatic reaction of r*Aae*UPO. Previously, the atmospheric pressure plasma jet (µAPPJ) was used for H_2_O_2_ generation in combination with r*Aae*UPO immobilized on amino-functionalized beads (ReliZyme HA403 M, Resindion) [13,20], resulting in a steady product formation. Due to a lack of plasma stability at high water vapor admixtures, it was only possible to enrich up to 25% of the feed gas with water. Since the H_2_O concentration greatly influences plasma-induced H_2_O_2_ generation [21], we tested the performance of the capillary plasma jet [26] in plasma-driven biocatalysis as it enables the use of higher input powers and thus the use of higher H_2_O concentrations in the feed gas (100% feed gas routed through the bubbler, 6400 ppm H_2_O). Both plasma sources were operated under conditions that allowed the highest H_2_O_2_ production and H_2_O_2_ accumulation in the liquid was analyzed (Figure 2). As previously observed for the microscale atmospheric pressure plasma jet [20], the H_2_O_2_ concentration increased linearly when the capillary plasma jet was operated with 100% gas being routed through the bubbler. However, the capillary jet showed a H_2_O_2_ production (based on the linear slope) of 0.14 mM min^-1^, while the microscale atmospheric pressure plasma jet produced 0.07 mM min^-1^. The increased H_2_O_2_ production of the capillary plasma jet presumably is caused by the increased amount of H_2_O in the feed gas: More water molecules can be dissociated by electron impact dissociation, and more H_2_O_2_ is generated via the recombination of OH radicals. Furthermore, the higher plasma power of the capillary plasma jet leads to increased electron density, which also leads to more dissociation by electron impact dissociation of water molecules [28].

**Figure 2:**
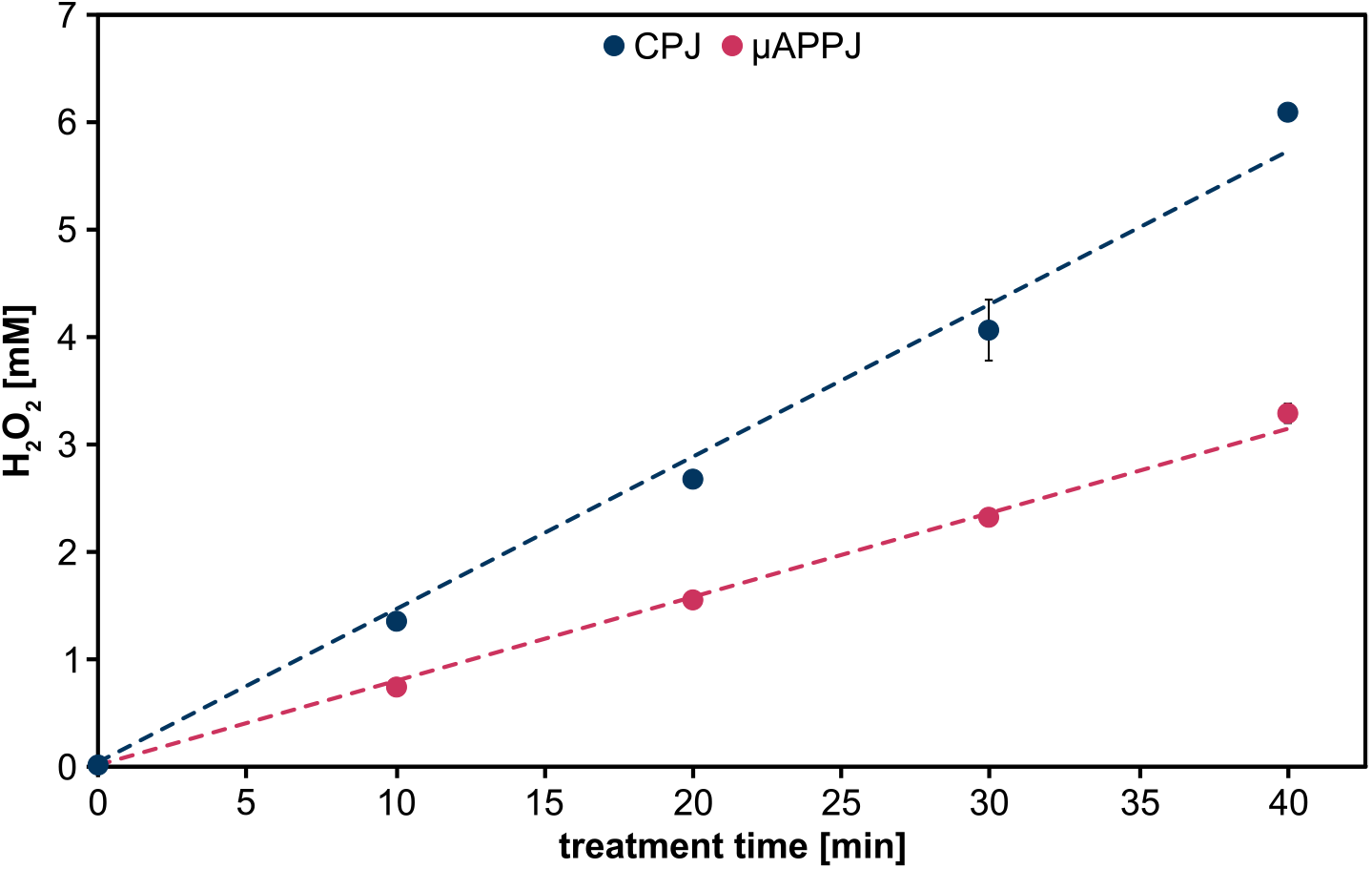
Time-dependent H_2_O_2_ production. Potassium phosphate buffer (5 ml, 100 mM, pH 7) was treated either with the capillary plasma jet (CPJ) and 6400 ppm water in the feed gas or the microscale atmospheric pressure plasma jet (µAPPJ) and 5750 ppm water in the feed gas for up to 40 min. The distances between the nozzles of the jets and the samples were approx. 16 mm (CPJ) and 6 mm (µAPPJ). Means and standard deviations reflect three experiments. Standard deviations that are not visible were below 0.1 mM.

In plasma-driven biocatalysis, the tailoring of the plasma and thus the amount of H_2_O_2_ supplied to match the needs and limitations of the enzyme appears very appealing, especially when employing highly H_2_O_2_-sensitive enzymes in the future. We therefore tested to what extent the H_2_O_2_ production by the capillary plasma jet can be influenced by different H_2_O concentrations in the feed gas (Figure 3). The range from 50 to 100% feed gas being passed through the bubbler had not been tested to prevent backflow of H_2_O to the mass flow controllers of the plasma source. The H_2_O_2_ concentration increased drastically upon introduction of water to the feed gas, showing the importance of H_2_O in the feed gas for H_2_O_2_ production in plasma-treated liquids. H_2_O_2_ increased from 0.13 mM (0 ppm H_2_O) to 0.37 mM (320 ppm H_2_O). In the absence of water addition to the gas flow the detected H_2_O_2_ presumably stems from water contamination. H_2_O_2_ formation is proportional to the H_2_O concentration in the feed gas, and the results are in good agreement with the previously reported measurements for the capillary plasma jet [28]. Doubling the H_2_O concentration from 320 ppm to 640 ppm led to an increase of H_2_O_2_ from 0.37 mM to 0.46 mM. Likely, this is attributable to a certain fraction of the formed H_2_O_2_ being degraded again in the gas phase. The maximum H_2_O_2_ concentration of 1.6 mM was reached at 10 min treatment time with 6400 ppm H_2_O in the feed gas, which equals 100% feed gas routed through the bubbler). This presents a great working range to adjust H_2_O_2_ production levels to biocatalyst’s needs.

**Figure 3:**
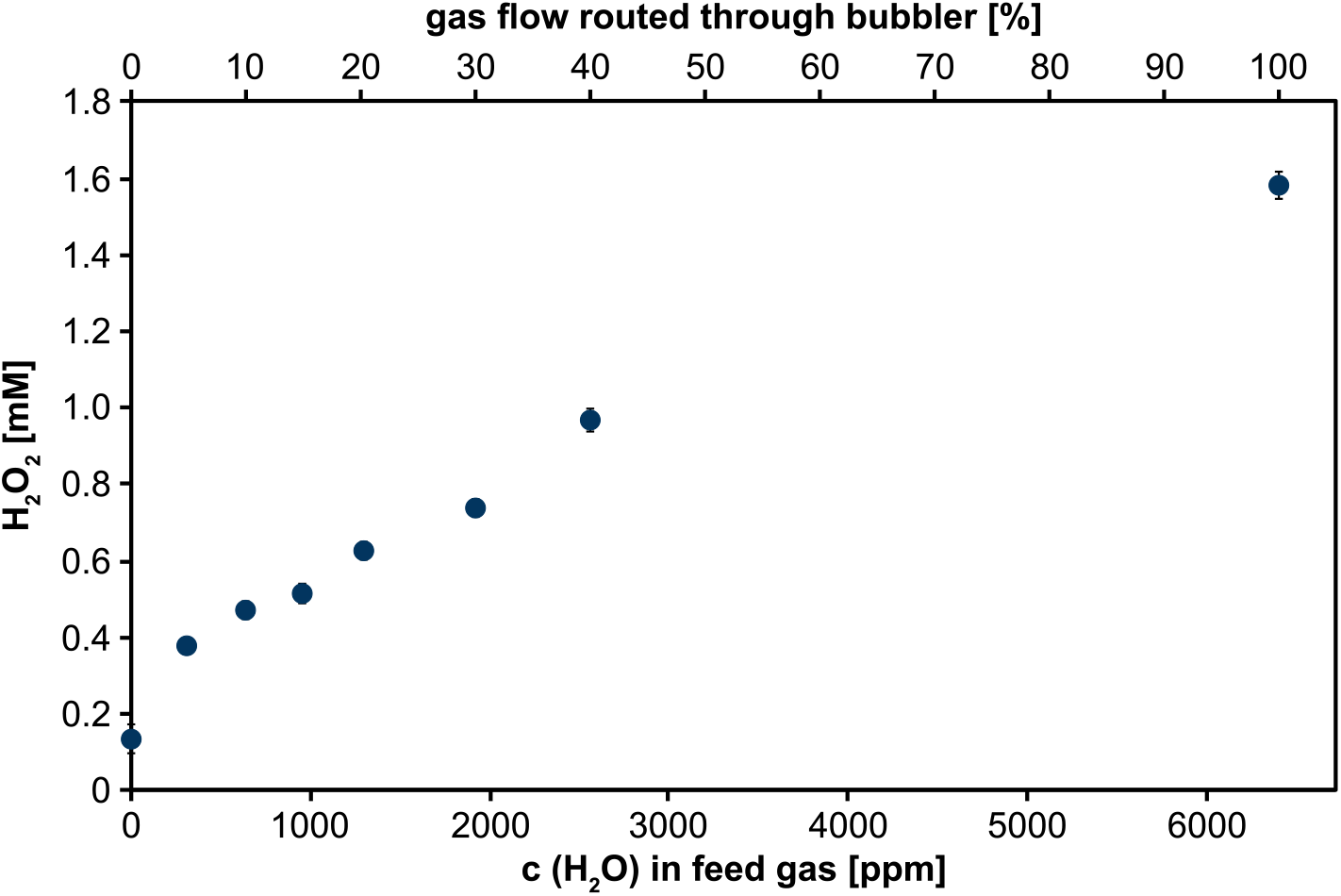
H_2_O_2_ production at different water concentrations in the feed gas. The feed gas of the capillary jet was (partially) routed through a cooled water-containing bubbler (in %) resulting in different concentrations of water (in ppm) in the feed gas. Potassium phosphate buffer (5 ml, 100 mM, pH 7) was treated for 10 min and the H_2_O_2_ concentration (in mM) was determined consecutively. Means and standard deviations represent three experiments shown. Standard deviations that are not visible were below 0.01 mM.

Using the capillary plasma jet with maximum water feed for plasma-driven biocatalysis, we tested the (*R*)-1-PhOl production using r*Aae*UPO immobilized on amino-functionalized ReliZyme beads (HA403 M) previously used for studying performance of the µAPPJ [20], as well as other ReliZyme beads (EA403 M, EP403 M, HFA403 M, BU403 M). Only for the HA304 M and the EA 403 M beads considerable product formation was observed (Figure 4, data for EP403 M, HFA403 M, BU403 M are displayed in Supplementary Figures 1-3). Product formation within the tested 40 min run time was comparable between both bead types (HA403 M and EA403 M) and reached a product concentration of approx. 1.7 mM (*R*)-1-PhOl (Figure 4). The turnover number for this system was 32,264 mol_(*R*)-1-PhOl_ mol^-1^_r*Aae*UPO_ for HA403 M and 25,261 mol_(*R*)-1-PhOl_ mol^-1^_r*Aae*UPO_ for EA403 M, which is in the same range as previously published results obtained using the µAPPJ and plasma-independent methods for H_2_O_2_ production (Supplementary Table 1 and 2) [20,30-32]. On both bead types enzyme inactivation was at 10% and thus residual activity was still very high (Supplementary Figure 4), indicating that the turnover number might be increased by prolonging the run time of the reaction. This experiment showed that not all beads are equally suited for plasma-driven biocatalysis with r*Aae*UPO.

**Figure 4:**
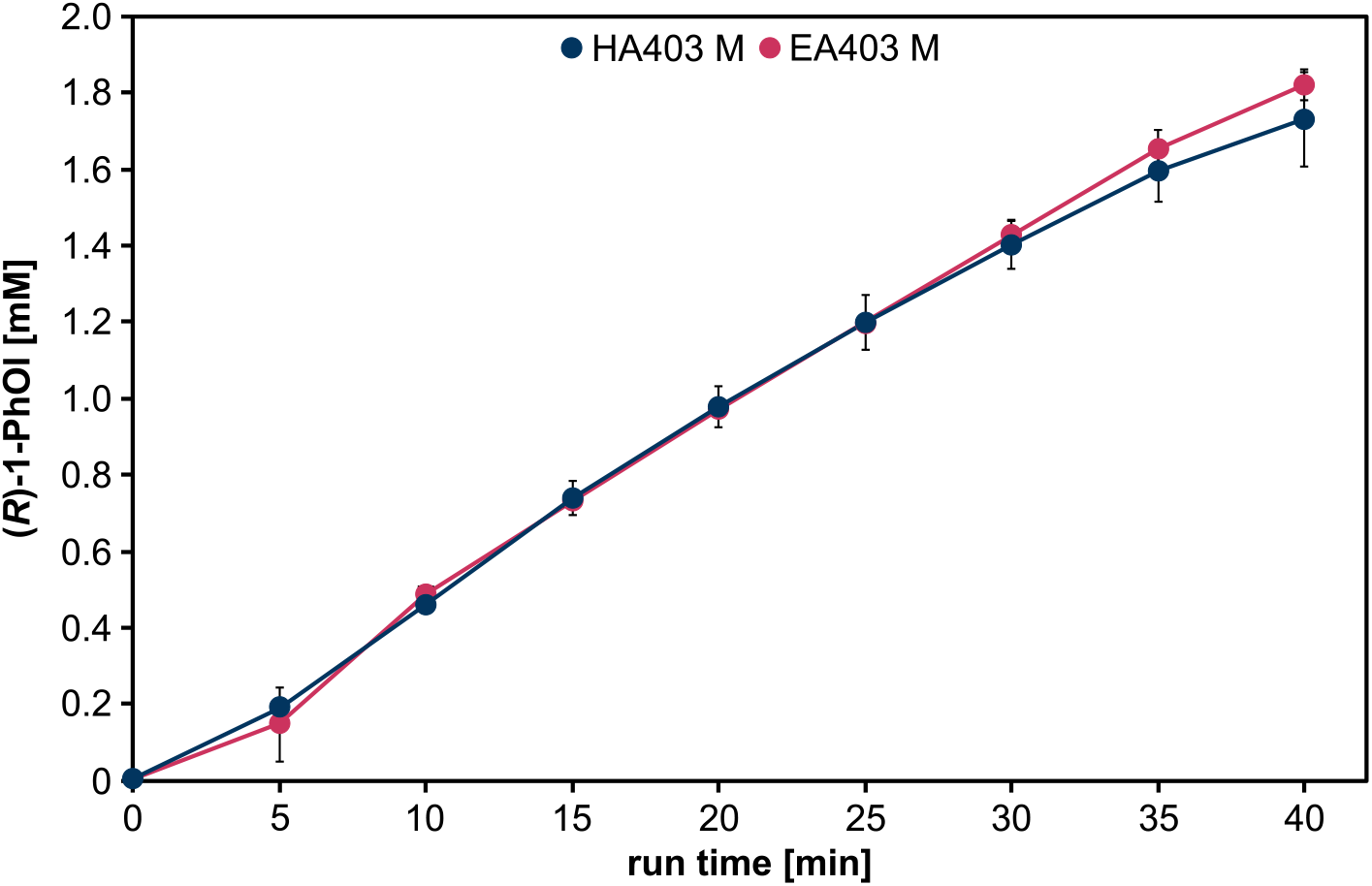
Plasma-driven biocatalysis with the capillary plasma jet using rAaeUPO immobilized on ReliZyme beads. Conversion of the substrate ETBE utilized H_2_O_2_ from direct plasma treatment of the r*Aae*UPO immobilized on ReliZyme HA403 M or EA403 M. Reaction solution contained 5 ml potassium phosphate buffer (100 mM, pH 7) with 50 mM ETBE. Plasma treatment was performed using the capillary plasma jet as described above, with a water concentration of 6400 ppm in the feed gas at 6 W plasma power. Every 5 min, aliquots were withdrawn for product analysis by GC. Means and standard deviations reflect three experiments.

In a previous study, r*Aae*UPO immobilized on resin-based carriers (Purolite) showed promising activity and plasma tolerance. Especially amino- and epoxy-butyl-functionalized beads showed a high level of plasma protection [18, 19]. The Purolite carriers were also tested for their potential use in plasma-driven biocatalysis with the capillary plasma jet (Figure 5). r*Aae*UPO immobilized on amino-functionalized resin ECR8309F showed comparable product formation to the enzyme on both ReliZyme beads, reaching a product concentration of 1.7 mM_(*R*)-1-PhOl_. The r*Aae*UPO on epoxy-butyl-functionalized resin ECR8285 only reached a product concentration of 1.0 mM_(*R*)-1-PhOl_. Using the Purolite beads, we were able to obtain turnover numbers of 32,708 mol_(*R*)-1-PhOl_ mol_-1r*Aae*UPO_ (ECR8309F) and 20,338 mol_(*R*)-1-PhOl_ mol_-1r*Aae*UPO_ (ECR8285), respectively. Maximum conversion rates were calculated from the initial linear slope of the reaction. They were similar for enzymes immobilized on either of the three amino-functionalized beads (approx. 50 µM_(*R*)-1-PhOl_ min^-1^), while the conversion rate for the enzyme on the epoxy-butyl beads was only 30 µM min^-1^ (Supplementary Table 5). The residual activity after 40 min biocatalysis was at of both Purolite carriers were decreased to 53% and 17% for ECR8309F and ECR8285, respectively, and thus lower than for the ReliZyme beads (Supplementary Figure 4). Weighting conversion rates and enzyme inactivation, the two ReliZyme bead types appear to be the more promising support materials under plasma-operating conditions.

**Figure 5:**
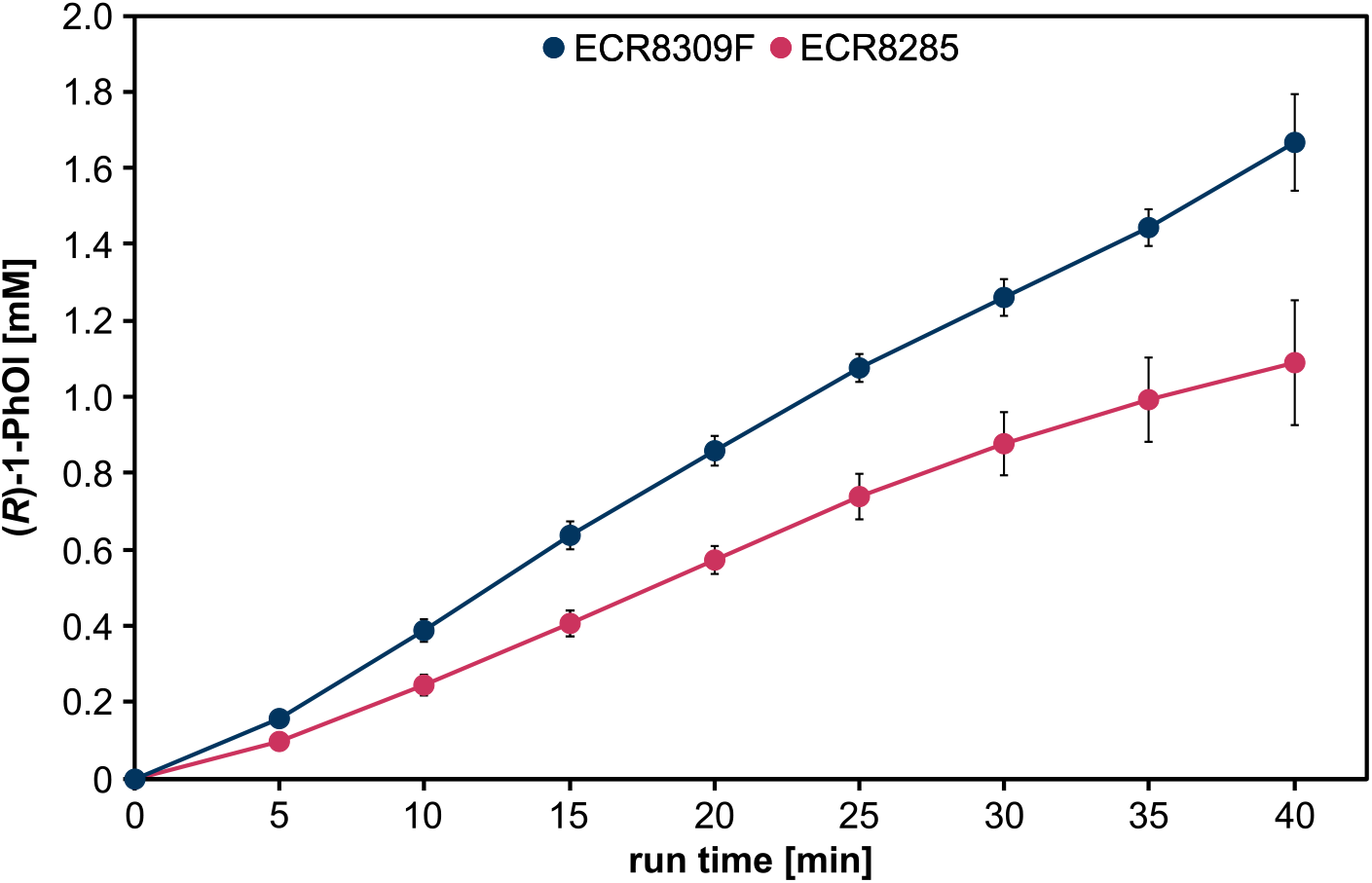
Plasma-driven biocatalysis with the capillary plasma jet and r*Aae*UPO immobilized on Purolite beads. The substrate ETBE was converted by r*Aae*UPO immobilized on Purolite ECR8309F and ECR8285. The reaction solution contained 5 ml potassium phosphate buffer (100 mM, pH 7) with 50 mM ETBE. H_2_O_2_ was generated by plasma treatment with the capillary plasma jet and a water concentration of 6400 ppm in the feed gas at 6 W plasma power. Every 5 min aliquots were withdrawn for product analysis by GC. Means and standard deviations represent three experiments.

Since for the ReliZyme beads enzyme inactivation after 40 min biocatalysis was negligible, the experiment was repeated increasing the run time to 80 min (Figure 6). After 40 min, for both bead types, product formation rates began to decrease, reaching a plateau at ∼2.3 mM_(*R*)-1-PhOl_ after 60 min. Investigation of the residual activity after biocatalysis revealed substantial enzyme inactivation (Supplementary Figure 5). Substrate conversion decelerated with increasing run times, resulting in H_2_O_2_ accumulation, which presumably is the main cause of enzyme inactivation in this system [5,6]. This notion is supported by the fact that after an 80 min biocatalysis with the µAPPJ [20], which produces only approx. 50% of the H_2_O_2_ produced by the capillary jet, the residual activity was still higher. Since residual enzyme activity was still at approx. 35-40% after 80 min biocatalysis using the capillary plasma jet, the turnover ceasing cannot be explained by enzyme inactivation alone. Rather, product inhibition and substrate limitation set in at this point, significantly inhibiting productivity the overall system [20]. Despite the reduced efficiency of biocatalysis during the prolonged run time, it was possible to increase the turnover number to 44,199 mol_(*R*)-1-PhOl_ mol_-1r*Aae*UPO_ (HA403 M) and 34,097 mol_(*R*)-1-PhOl_ mol_-1r*Aae*UPO_ (EA403 M) using the capillary plasma jet, respectively (Supplementary Table 1 and 2).

**Figure 6:**
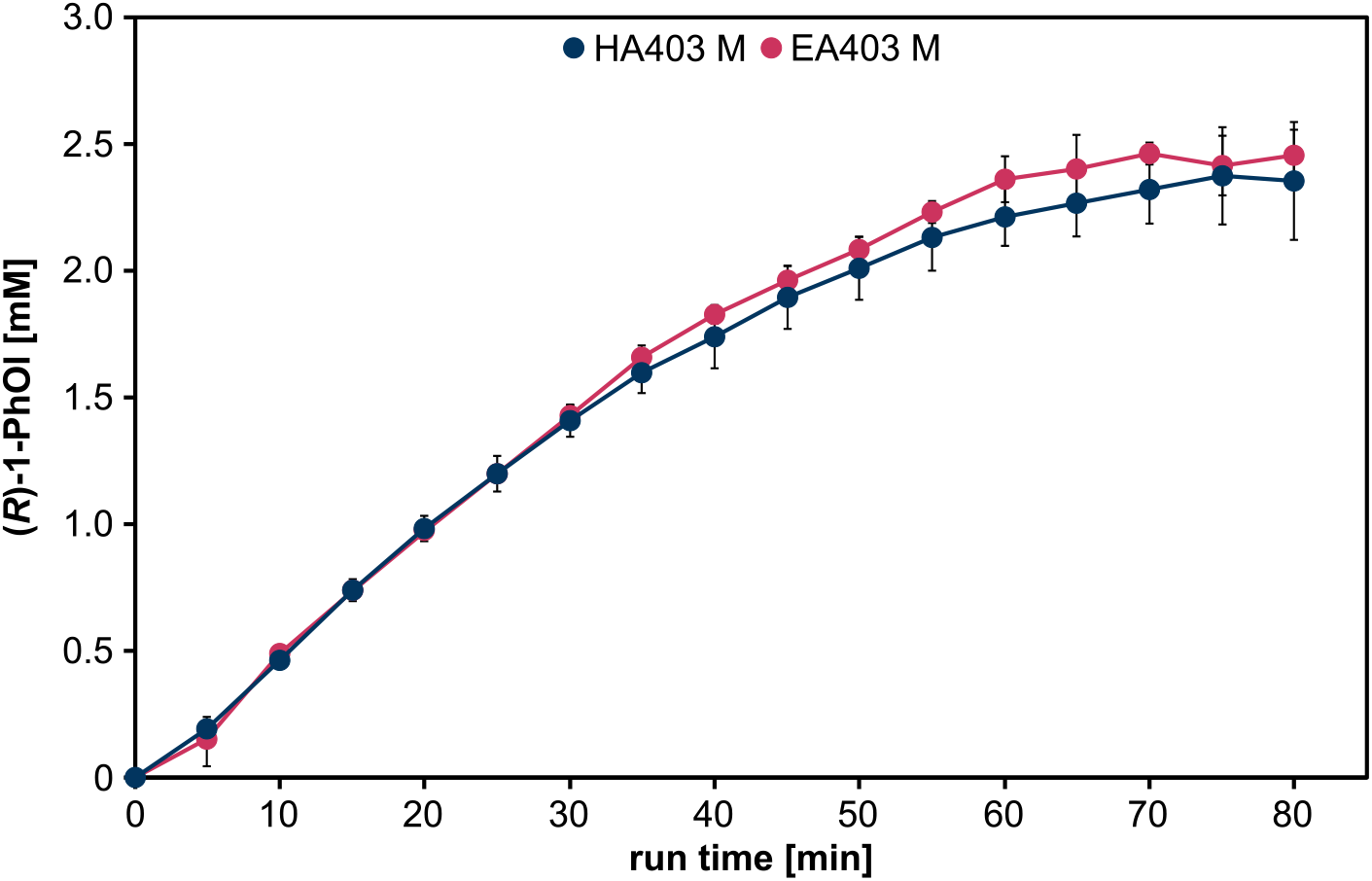
Prolonged plasma-driven biocatalysis with capillary plasma jet. Conversion of the substrate ETBE and H_2_O_2_ from plasma treatment by r*Aae*UPO immobilized on ReliZyme HA403 M and EA403 M was performed as described above, with the capillary plasma jet and a water concentration of 6400 ppm in the feed gas at 6 W plasma power. Every 5 min aliquots were withdrawn for product quantification by GC. Means and standard deviations reflect three experiments.

For plasma-driven biocatalysis turnover number achieved with r*Aae*UPO immobilized on HA403 M beads and replacing the entire buffer system every 10 min was 36,414 mol_(*R*)-1-PhOl_ mol^-1^_r*Aae*UPO_ [20]. In the present study, the same efficiency of plasma-driven biocatalysis was reached without having to replace the reaction solution.

To circumvent the limitations of product inhibition and substrate limitation, a long-term experiment was carried out in which the r*Aae*UPO was immobilized on HA403 M beads and the reaction solution was replaced every 5 or 10 min (Figure 7). At a product formation per minute below 0.05 µmol min^-1^ the enzyme was considered inactive and the reaction was stopped. Using a water vapor concentration of 6400 ppm combined with a reaction solution exchange every 10 min, product formation rates between 0.25 and 0.3 µmol (*R*)-1-Phol min^-1^ were observed in the first two hours of biocatalysis.

**Figure 7:**
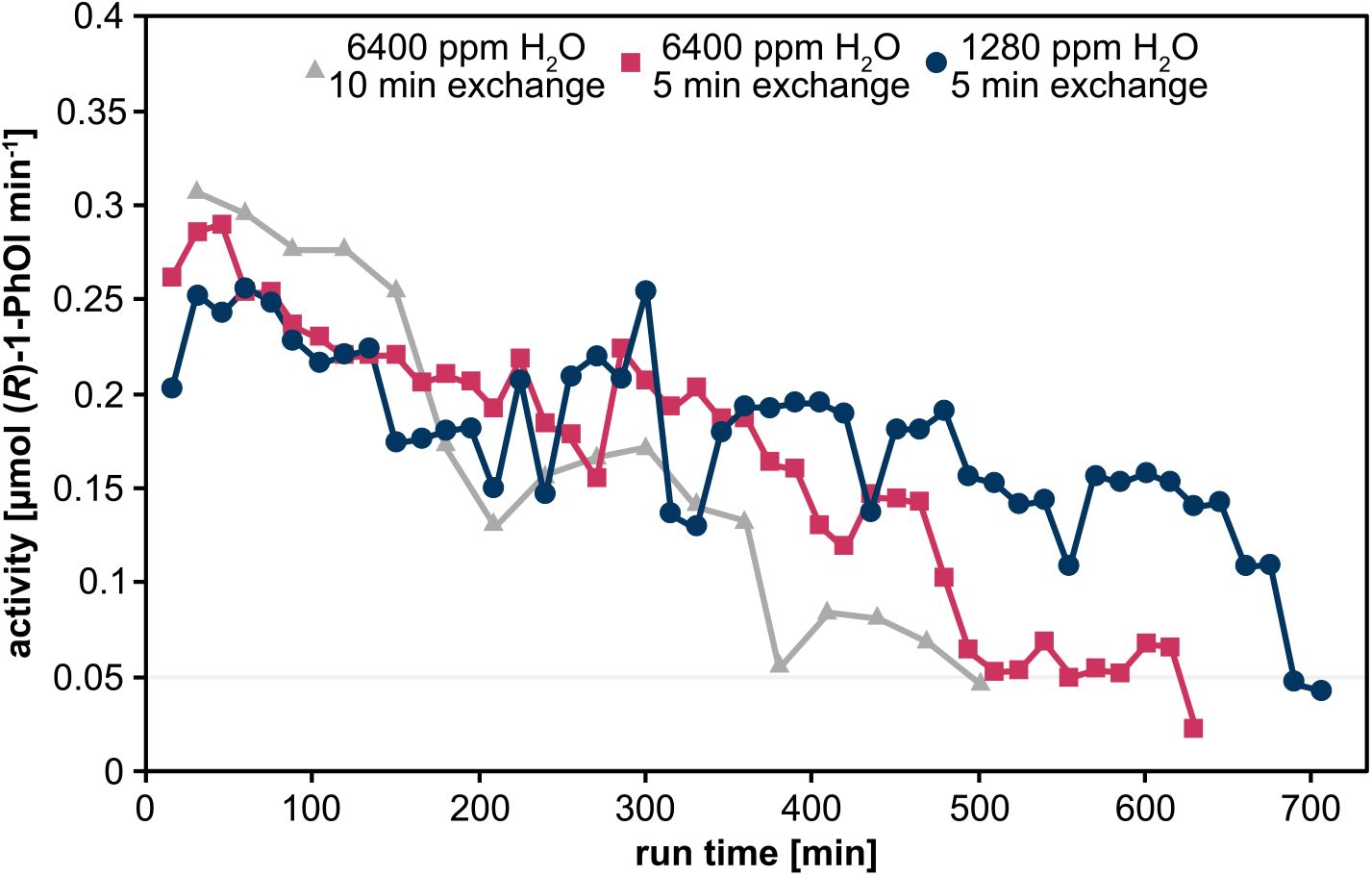
Product formation per minute in long-term experiments with frequent exchange of reaction solutions. Activity of r*Aae*UPO immobilized on HA403 M beads is shown as a function of plasma-driven biocatalysis run time. Every 10 or 5 min, the complete reaction solution was exchanged, and product formation was analyzed by GC measurement. Either 1280 ppm or 6400 ppm H_2_O were added to the feed gas to modulate H_2_O_2_ generation. Means reflect two experiments. A detailed presentation including standard deviations for each experimental condition is provided in the supplements (Supplementary Figures 6-8).

Afterwards, activity decreased rapidly to 0.15 µmol min^-1^ after 200 min and 0.08 µmol min^-1^ after 400 min, and the biocatalytic reaction was stopped after 500 min run time. Based on the H_2_O_2_ generation data shown in Figure 3, it is known that using a water concentration of 6400 ppm in the feed gas results in the generation of approx. 1.6 mM H_2_O_2_ in 10 min, which corresponds to 0.8 µmol H_2_O_2_ min^-1^. Thus, at the maximum conversion rate of the enzyme, only half of the generated H_2_O_2_ was converted into (*R*)-1-PhOl. This likely resulted in an excess of H_2_O_2_ (theoretically of several µmol) within 10 min incubation in one buffer solution. Thus, we assume that the rapid loss of activity after 120 min run time is due to H_2_O_2_-induced inactivation of r*Aae*UPO. To minimize enzyme incubation in H_2_O_2_-excess conditions, we reduced the time for the exchange of the reaction solution to 5 min. This resulted in a stabilized production range of 0.17 to 0.25 µmol (*R*)-1-Phol per cycle for 460 min. For the 5 min exchange condition, reaction was stopped after 630 min. Thus, stable enzyme performance was extended from 120 to 480 min. Since a more frequent buffer exchange positively affected r*Aae*UPO activity, the experiment was repeated with less H_2_O in the feed gas to limit the potential H_2_O_2_ excess. Based on the maximum conversion rate of ethylbenzene of 0.3 µmol min^-1^, a water vapor admixture of 1280 ppm to the feed gas was chosen, 0.3 µmol min^-1^ H_2_O_2_ (see Figure 3). Product formation under these conditions averaged about 0.2 µmol min^-1^ (Figure 7). Activity of the r*Aae*UPO remained above 0.1 µmol (*R*)-1-Phol min^-1^ until a treatment time of 675 min was reached. The reaction was stopped after 705 min. Inactivation is likely still attributable to an H_2_O_2_ excess, since not all the 0.3 µmol H_2_O_2_ min^-1^ generated in the cycle were converted and further reduction in H_2_O_2_ could positively affect enzyme lifetime of r*Aae*UPO. This highlights that tailoring of plasma conditions can go a long way to accommodating to the requirements of the enzyme.

In addition to activity over time, the total amount of product accumulated over time was analyzed (Figure 8). The fastest product increase was observed when 6400 ppm H_2_O were admixed and reaction solution was exchanged every 10 min. Reflecting enzyme inactivation over time, the product accumulation rates declined after approx. 150 min, and after 230 min. The total amount of product generated was lower compared to the more frequent reaction solution exchange, and the admixture of only 1280 ppm H_2_O combined with 5 min cycles, respectively. The use of 1280 ppm H_2_O in the feed gas led to the slowest increase in product but gave the highest final yield of 122 µmol_(*R*)-1-PhOl_, compared to 84 µmol at 6400 ppm H_2_O and 10 min cycles and 102 µmol at 6400 ppm H_2_O and 5 min cycles. Accordingly, the highest total turnover number was observed for 1280 ppm H_2_O in the feed gas and 5 min cycle times, which was a TTN of 174,209 mol_(*R*)-1-PhOl_ mol^-1^_r*Aae*UPO_ (Table 1, detailed list for each condition is given in Supplementary tables 6-8).

**Table 1:** Total product formation and TTN calculations of long-term experiment.

| Method | Total product [μmol] | TTN<br>[mol<sub>(R)-1-PhOI</sub> mol<sup>-1</sup><sub>rAaeUPO</sub>] |
| --- | --- | --- |
| 6400 ppm H<sub>2</sub>O, 10 min buffer exchange | 84 ± 12 | 122,138 ± 5,694 |
| 6400 ppm H<sub>2</sub>O, 5 min buffer exchange | 102 ± 10 | 138,777 ± 22,687 |
| 1280 ppm H<sub>2</sub>O, 5 min buffer exchange | 122 ± 4 | 174,209 ± 3,921 |

**Figure 8:**
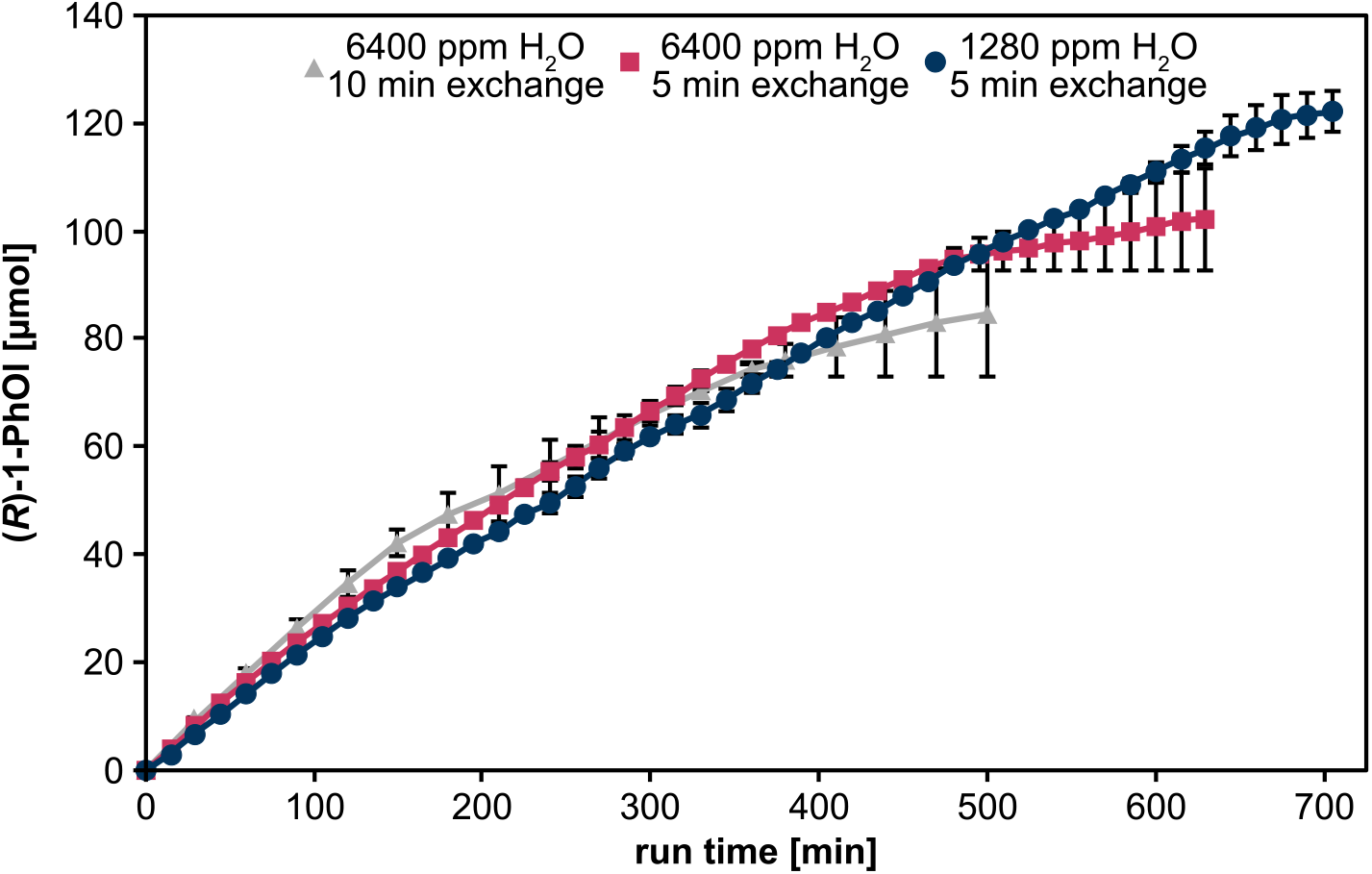
Total product accumulation in the long-term experiment over 5 or 10 min cycles. Accumulation of generated product (*R*)-1-PhOl produced by r*Aae*UPO immobilized on HA403 M beads after different plasma-driven biocatalysis run times. Either 6400 ppm or 1280 ppm H_2_O were added to the feed gas for H_2_O_2_ generation. Every 10 or 5 min, the complete reaction solution was exchanged, and product formation was analyzed by GC measurement Means and standard deviations representing two experiments are shown. Standard deviations that are not visible were below 2 µmol.

In summary, the turnover number of plasma-driven biocatalysis was increased almost five-fold through optimization of H_2_O_2_ production operating the capillary plasma jet with different H_2_O admixtures and extending the run time of the process. With a total turnover number of 174,209 mol_(*R*)-1-PhOl_ mol^-1^_r*Aae*UPO_, the process of plasma-driven biocatalysis is in the upper range of previously reported total turnover numbers of *Aae*UPO-mediated conversion of ETBE with H_2_O_2_ generated by *in situ* methods [7,30-32]. While the efficiency of the process can be further optimized by matching enzyme operating parameters and H_2_O_2_ even more closely and by adjusting the reactor design, these results highlight that plasma-driven biocatalysis is a competitive method when it comes to enzyme-compatible methods of *in situ* H_2_O_2_ generation.

## Author’s contributions

T.D.: conceptualization, formal analysis, investigation, visualization, writing of original draft; D.S.: investigation; S.S.: resources, writing – review and editing; F.H.: resources, writing – review and editing; J.G.: resources, writing – review and editing; J.E.B.: conceptualization, funding acquisition, supervision, writing – review and editing.

