## Supplementary Information for "The atmospheric pressure capillary plasma jet is well-suited to supply H_2_O_2_ for plasma-driven biocatalysis"

**Supplementary Table 1: Calculations of r*Aae*UPO concentrations and TON using HA403 M beads and the capillary plasma jet with 6400 ppm $H_2O$ in feed gas.**

| | |
|---|---|
| **Enzyme loading of beads in immobilization** | |
| amount of beads [mg] | 500 |
| total volume [ml] | 5 |
| r*Aae*UPO concentration [nM] | 400 |
| r*Aae*UPO amount [nmol] | 2 |
| maximum loading of beads [nmol/100 mg beads] | 0.4 |
| binding efficiency [%] | 44.68 |
| actual loading of beads [nmol/100 mg] | 0.18 |
| **Final concentrations of r*Aae*UPO in reaction** | |
| volume used for reactor [ml] | 1.5 |
| amount of beads in reactor [mg] | 150 |
| amount r*Aae*UPO in reactor [nmol] | 0.27 |
| reaction volume [ml] | 5 |
| r*Aae*UPO concentration in reaction [nM] | 53.62 |
| **TON calculations after 40 min** | |
| product concentration [mM] | 1.73 |
| turnover number | 32,265 |
| **TON calculations after 80 min** | |
| product concentration [mM] | 2.37 |
| turnover number | 44,200 |

**Supplementary Table 2: Calculations of r*Aae*UPO concentrations and TON using EA403 M beads the capillary plasma jet with 6400 ppm $H_2O$ in feed gas.**

| **Enzyme loading of beads in immobilization** | |
|---|---|
| amount of beads [mg] | 500 |
| total volume [ml] | 5 |
| r*Aae*UPO concentration [nM] | 400 |
| r*Aae*UPO amount [nmol] | 2 |
| maximum loading of beads [nmol/100 mg beads] | 0.4 |
| binding efficiency [%] | 59.95 |
| actual loading of beads [nmol/100 mg] | 0.24 |
| **Final concentrations of r*Aae*UPO in reaction** | |
| volume used for reactor [ml] | 1.5 |
| amount of beads in reactor [mg] | 150 |
| amount r*Aae*UPO in reactor [nmol] | 0.36 |
| reaction volume [ml] | 5 |
| r*Aae*UPO concentration in reaction [nM] | 71.95 |
| **TON calculations after 40 min** | |
| product concentration [mM] | 1.81 |
| turnover number | 25,261 |
| **TON calculations after 80 min** | |
| product concentration [mM] | 2.45 |
| turnover number | 34,097 |

**Supplementary Table 3: Calculations of r*Aae*UPO concentrations and TON using ECR8309F beads the capillary plasma jet with 6400 ppm $H_2O$ in feed gas.**

| **Enzyme loading of beads in immobilization** | |
|---|---|
| amount of beads [mg] | 500 |
| total volume [ml] | 5 |
| r*Aae*UPO concentration [nM] | 400 |
| r*Aae*UPO amount [nmol] | 2 |
| maximum loading of beads [nmol/100 mg beads] | 0.4 |
| binding efficiency [%] | 42.33 |
| actual loading of beads [nmol/100 mg] | 0.17 |
| **Final concentrations of r*Aae*UPO in reaction** | |
| volume used for reactor [ml] | 1.5 |
| amount of beads in reactor [mg] | 150 |
| amount r*Aae*UPO in reactor [nmol] | 0.25 |
| reaction volume [ml] | 5 |
| r*Aae*UPO concentration in reaction [nM] | 50.80 |
| **TON calculations after 40 min** | |
| product concentration [mM] | 1.66 |
| turnover number | 32,708 |

**Supplementary Table 4: Calculations of r*Aae*UPO concentrations and TON using ECR8285 beads the capillary plasma jet with 6400 ppm $H_2O$ in feed gas.**

| **Enzyme loading of beads in immobilization** | |
|---|---|
| amount of beads [mg] | 500 |
| total volume [ml] | 5 |
| r*Aae*UPO concentration [nM] | 400 |
| r*Aae*UPO amount [nmol] | 2 |
| maximum loading of beads [nmol/100 mg beads] | 0.4 |
| binding efficiency [%] | 44.40 |
| actual loading of beads [nmol/100 mg] | 0.18 |
| **Final concentrations of r*Aae*UPO in reaction** | |
| volume used for reactor [ml] | 1.5 |
| amount of beads in reactor [mg] | 150 |
| amount r*Aae*UPO in reactor [nmol] | 0.27 |
| reaction volume [ml] | 5 |
| r*Aae*UPO concentration in reaction [nM] | 53.28 |
| **TON calculations after 40 min** | |
| product concentration [mM] | 1.08 |
| turnover number | 20,338 |

**Supplementary Table 5: Calculations of the linear slope of r*Aae*UPO in plasma-driven biocatalysis using different carriers for immobilization and the capillary plasma jet with 6400 ppm $H_2O$ in feed gas.**

| **Linear slope [µM (*R*)-1-PhOl $min^{-1}$]** | |
|---|---|
| HA403 M | 49.03 |
| EA403 M | 54.57 |
| ECR8309F | 47.32 |
| ECR8285 | 31.90 |

**Supplementary Table 6: Calculations of TTN using r*Aae*UPO immobilized on HA403 M beads in long-term biocatalysis with the capillary plasma jet.** 6400 ppm $H_2O$ was added to the feed gas and buffer exchange was performed every 10 min.

| | |
|---|---|
| **Enzyme loading of beads in immobilization** | |
| amount of beads [mg] | 500 |
| total volume [ml] | 5 |
| r*Aae*UPO concentration [nM] | 400 |
| r*Aae*UPO amount [nmol] | 2 |
| maximum loading of beads [nmol/100 mg beads] | 0.4 |
| binding efficiency [%] | 43.00 |
| actual loading of beads [nmol/100 mg] | 0.17 |
| **Final concentrations of r*Aae*UPO in reaction** | |
| volume used for reactor [ml] | 4 |
| amount of beads in reactor [mg] | 400 |
| amount r*Aae*UPO in reactor [nmol] | 0.69 |
| reaction volume [ml] | 5 |
| r*Aae*UPO concentration in reaction [nM] | 137.61 |
| **TTN calculation** | |
| product [µmol] | 83.43 |
| total turnover number | 122,138 |

**Supplementary Table 7: Calculations of TTN using r*Aae*UPO immobilized on HA403 M beads in long-term biocatalysis with the capillary plasma jet.** 6400 ppm $H_2O$ was added to the feed gas and buffer exchange was performed every 5 min.

| | |
|---|---|
| **Enzyme loading of beads in immobilization** | |
| amount of beads [mg] | 500 |
| total volume [ml] | 5 |
| r*Aae*UPO concentration [nM] | 400 |
| r*Aae*UPO amount [nmol] | 2 |
| maximum loading of beads [nmol/100 mg beads] | 0.4 |
| binding efficiency [%] | 47.14 |
| actual loading of beads [nmol/100 mg] | 0.19 |
| **Final concentrations of r*Aae*UPO in reaction** | |
| volume used for reactor [ml] | 4 |
| amount of beads in reactor [mg] | 400 |
| amount r*Aae*UPO in reactor [nmol] | 0.75 |
| reaction volume [ml] | 5 |
| r*Aae*UPO concentration in reaction [nM] | 150.85 |
| **TTN calculations** | |
| product [µmol] | 102.13 |
| total turnover number | 138,777 |

**Supplementary Table 8: Calculations of TTN using r*Aae*UPO immobilized on HA403 M beads in long-term biocatalysis with the capillary plasma jet.** 1280 ppm $H_2O$ was added to the feed gas and buffer exchange was performed every 5 min.

| **Enzyme loading of beads in immobilization** | |
|---|---|
| amount of beads [mg] | 500 |
| total volume [ml] | 5 |
| r*Aae*UPO concentration [nM] | 400 |
| r*Aae*UPO amount [nmol] | 2 |
| maximum loading of beads [nmol/100 mg beads] | 0.4 |
| binding efficiency [%] | 43.85 |
| actual loading of beads [nmol/100 mg] | 0.17 |
| **Final concentrations of r*Aae*UPO in reaction** | |
| volume used for reactor [ml] | 4 |
| amount of beads in reactor [mg] | 400 |
| amount r*Aae*UPO in reactor [nmol] | 0.70 |
| reaction volume [ml] | 5 |
| r*Aae*UPO concentration in reaction [nM] | 140.32 |
| **TTN calculations** | |
| product [μmol] | 122.06 |
| total turnover number | 174,209 |

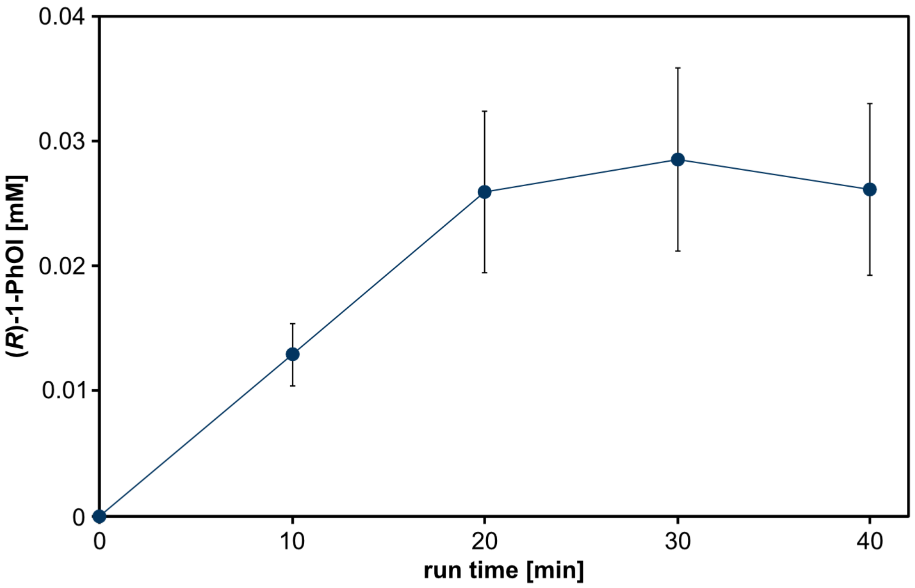


**Supplementary Figure 1: Plasma-driven biocatalysis with the capillary plasma jet using rAaeUPO immobilized on ReliZyme EP403 M beads.** Conversion of the substrate ETBE utilized $H_2O_2$ from direct plasma treatment of the r*Aae*UPO immobilized on ReliZyme EP403 M. Reaction solution contained 5 ml potassium phosphate buffer (100 mM, pH 7) with 50 mM ETBE. Plasma treatment was performed using the capillary plasma jet as described above, with a water concentration of 6400 ppm in the feed gas at 6 W plasma power. Every 10 min, aliquots were withdrawn for product analysis by GC. Means and standard deviations reflect three experiments.

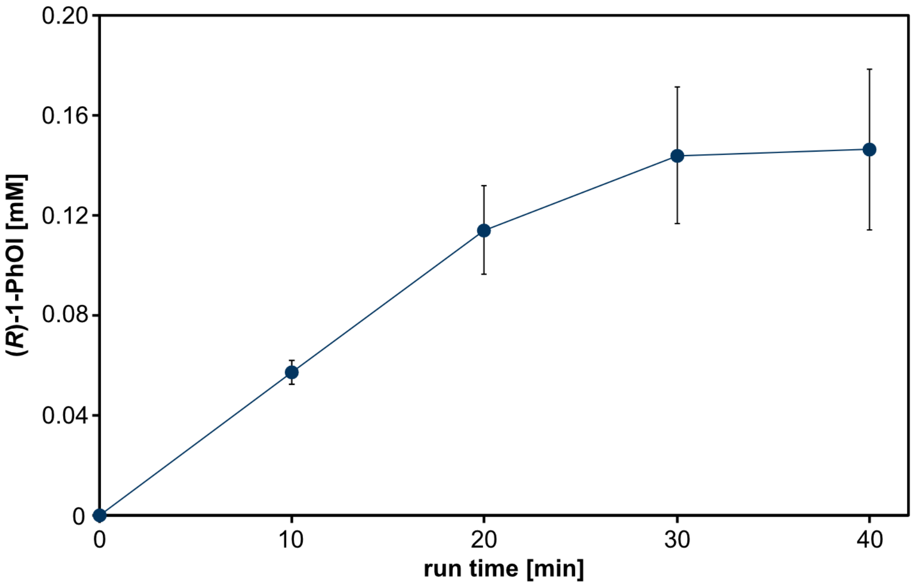


**Supplementary Figure 2: Plasma-driven biocatalysis with the capillary plasma jet using rAaeUPO immobilized on ReliZyme HFA403 M beads.** Conversion of the substrate ETBE utilized $H_2O_2$ from direct plasma treatment of the r*Aae*UPO immobilized on ReliZyme HFA403 M. Reaction solution contained 5 ml potassium phosphate buffer (100 mM, pH 7) with 50 mM ETBE. Plasma treatment was performed using the capillary plasma jet as described above, with a water concentration of 6400 ppm in the feed gas at 6 W plasma power. Every 10 min, aliquots were withdrawn for product analysis by GC. Means and standard deviations represent three experiments.

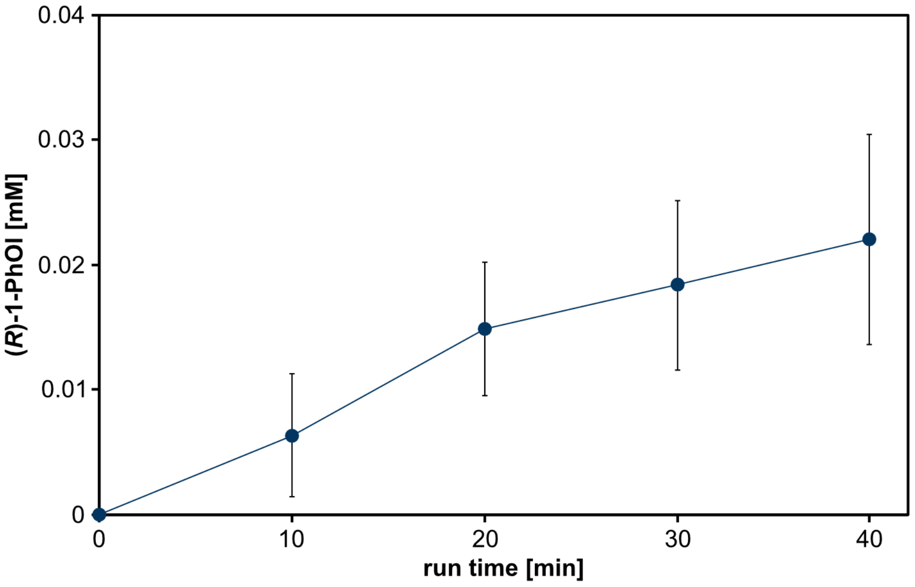


**Supplementary Figure 3: Plasma-driven biocatalysis with the capillary plasma jet using rAaeUPO immobilized on ReliZyme BU403 M beads.** Conversion of the substrate ETBE utilized $H_2O_2$ from direct plasma treatment of the r*Aae*UPO immobilized on ReliZyme BU403 M. Reaction solution contained 5 ml potassium phosphate buffer (100 mM, pH 7) with 50 mM ETBE. Plasma treatment was performed using the capillary plasma jet as described above, with a water concentration of 6400 ppm in the feed gas at 6 W plasma power. Every 10 min, aliquots were withdrawn for product analysis by GC. Means and standard deviations reflect three experiments.

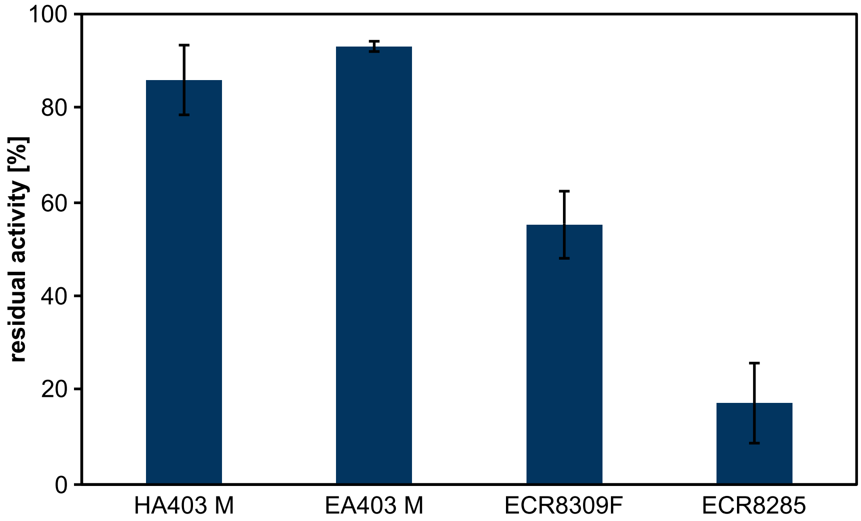


**Supplementary Figure 4: Residual activity of r*Aae*UPO on ReliZyme (HA403 M, EA403 M) and Purolite (ECR8309F and ECR8285) beads after 40 min plasma-driven biocatalysis.** After biocatalysis, enzyme-loaded beads were recovered and washed thrice with potassium phosphate buffer (100 mM, pH 7). Enzyme activity was determined using 2.5 mM 2,2'-azino-bis(3-ethylbenzothiazoline-6-sulfonic acid) (ABTS), 1 mM $H_2O_2$ and 50 mM citrate. Samples were shaken during turnover to ensure sufficient substrate supply. Every two minutes in a total of ten minutes reaction time, aliquots of 100 µl were withdrawn and measured at 405 nm using a microplate reader (Biotek Epoch). Enzyme activity was calculated based on the linear slope of the kinetic. Means and standard deviations represent three experiments.

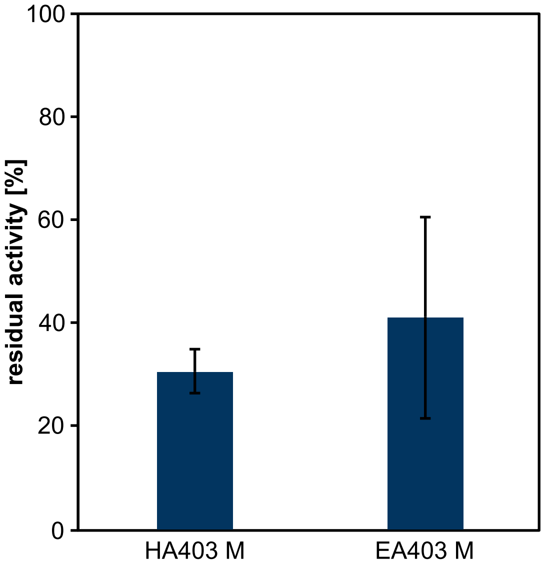


**Supplementary Figure 5: Residual activities of r*Aae*UPO on ReliZyme beads after 80 min plasma-driven biocatalysis with the capillary plasma jet.** Biocatalysis was performed with the capillary plasma jet and a water concentration of 6400 ppm in the feed gas at 6 W plasma power for a run time of 80 min. After biocatalysis, enzyme-loaded beads were recovered and washed thrice with potassium phosphate buffer (100 mM, pH 7). Enzyme activity was determined using 2.5 mM 2,2'-azino-bis(3-ethylbenzothiazoline-6-sulfonic acid) (ABTS), 1 mM $H_2O_2$ and 50 mM citrate. Samples were shaken during turnover to ensure sufficient substrate supply. Every two minutes in a total of ten minutes reaction time, aliquots of 100 µl were removed and measured at 405 nm using a microplate reader (Biotek Epoch). Enzyme activity was calculated based on the linear slope of the kinetic. Means and standard deviations reflect three experiments.

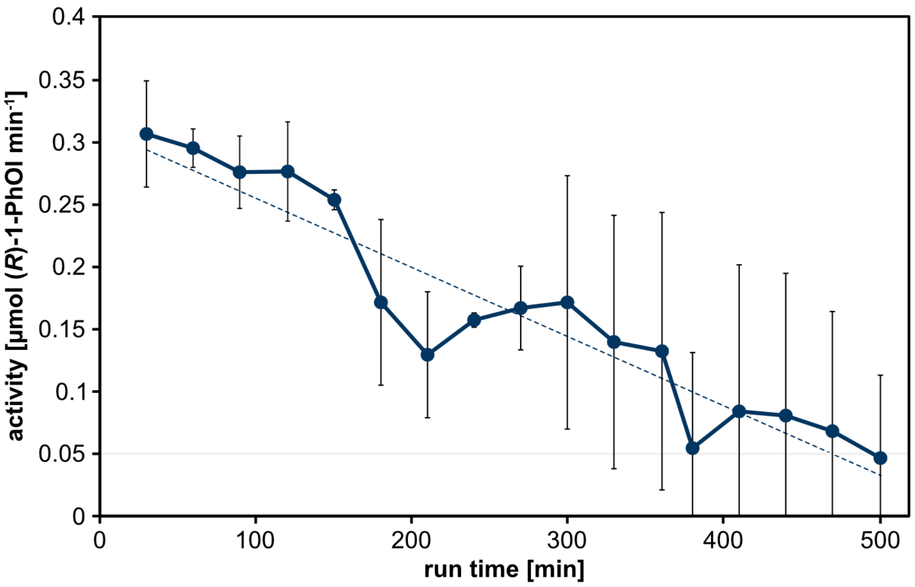


**Supplementary Figure 6: Product formation per minute in long-term experiments using 6400 ppm $H_2O$ in the feed gas.** Activity of r*Aae*UPO immobilized on HA403 M beads is plotted as a function of plasma-driven biocatalysis run time. Every 10 min, the complete reaction solution was exchanged and product formation was analyzed by GC measurement. Means and standard deviations reflect three experiments.

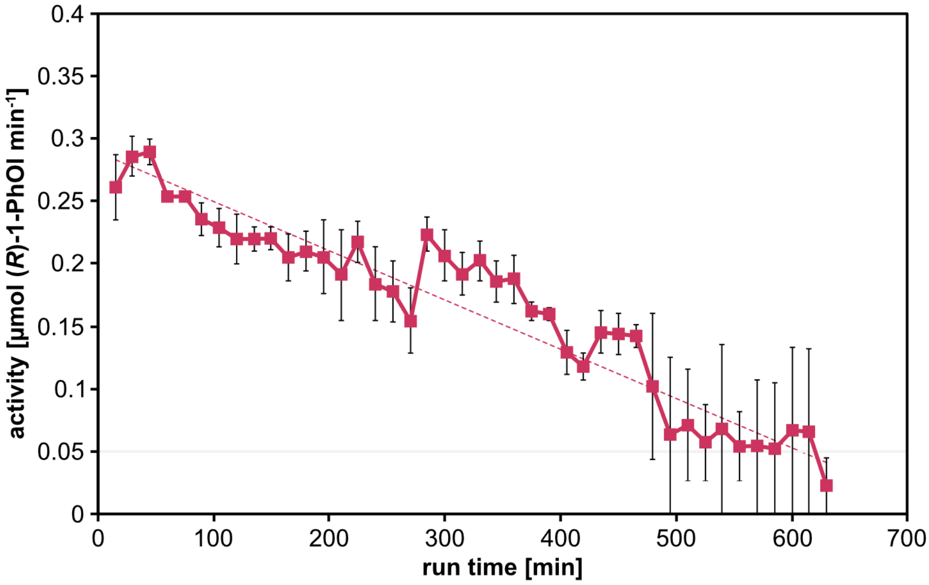


**Supplementary Figure 7: Product formation per minute in long-term experiments using 6400 ppm $H_2O$ in the feed gas.** Activity of r*Aae*UPO immobilized on HA403 M beads is plotted as a function of plasma-driven biocatalysis run time. Every 5 min, the complete reaction solution was exchanged and product formation was analyzed by GC measurement. Means and standard deviations reflect three experiments.

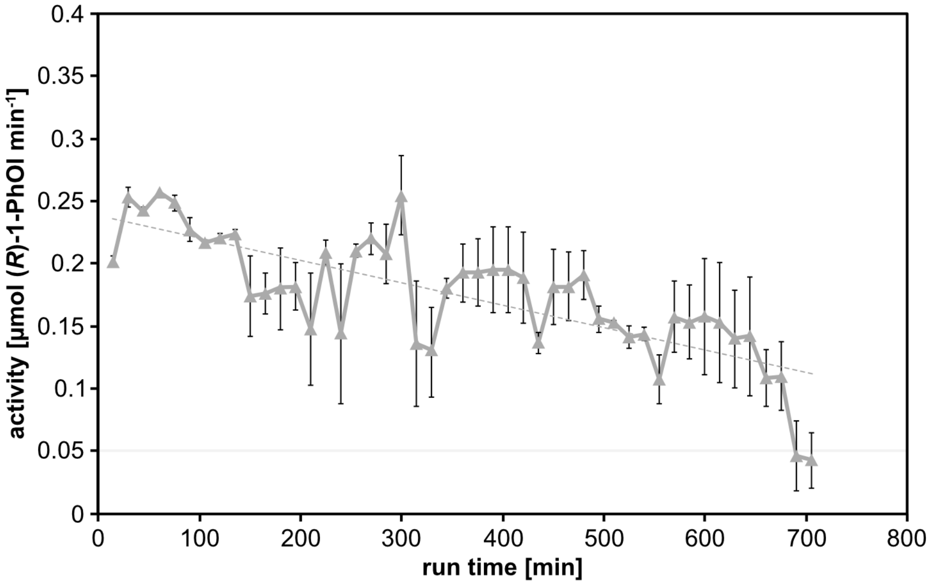


**Supplementary Figure 8: Product formation per minute in long-term experiments using 1280 ppm $H_2O$ in the feed gas.** Activity of r*Aae*UPO immobilized on HA403 M beads is plotted as a function of plasma-driven biocatalysis run time. Every 5 min, the complete reaction solution was exchanged and product formation was analyzed by GC measurement. Means and standard deviations represent three experiments.
